# Erythrocyte indices, anaemia levels and types in Kenyan injection and non-injection substance users

**DOI:** 10.1101/434365

**Authors:** Emmanuel Mulaya Khazalwa, Tom Were, David Hughes Mulama, Valentine Budambula

## Abstract

The impact of injection and non-injection substance use in HIV infections is an area of great public importance especially with respect to hematologic and immune profiles. Evaluations of hematologic and immune status are critical for better disease classification and clinical management especially of HIV positive substance users. However, not much information is known about the hematologic and immune derangements in HIV infected injection and non-injection substance users. This study, therefore, aimed at determining the laboratory markers of hematologic and immune derangements in HIV infected substance users. Hematologic and immune profiles were evaluated on venous blood specimens obtained from injection substance users, ISU (HIV-infected, n=62 and -uninfected, n=213) and non-injection substance users (HIV-infected, n=33 and -uninfected, n=186); and non-substance using controls (n=56) from Mombasa, coastal town of Kenya. The prevalence of anemia was higher in HIV infected ISU (48.4%) and non-ISU (63.6%) (p<0.0001); and HIV uninfected ISUs (56.3%) compared to HIV-uninfected non-ISUs (39.2%) and non-substance using controls (28.6%; p=0.0028). Hypochromic anaemia was more prevalent in the HIV-infected ISU (50.0%) and non-ISU (61.9%), and HIV-negative ISU (63.3%) relative to the HIV-negative non-ISU (39.7%) and non-substance using controls (56.3%; p=0.0007). Mild immunodeficiency dominated in the HIV infected individuals (HIV-infected ISU, 32.3% and non-ISU, 21.2%) versus HIV-uninfected ISU (16.9%); non-ISU (12.9%); and non-substance users (14.3%) while severe immunosuppression prevailed in HIV infected substance users (ISU, 14.5% and non-ISU, 15.2%) against HIV uninfected substance users (ISU, 5.2% and non-ISU, 3.8%); thus immunosuppression in substance users is aggravated with HIV infection. Moreover, drug-induced immunosuppression is associated with a higher likelihood of anaemia in HIV-uninfected substance users; ISU (OR=3.95, CI=1.934-8.077, p<0.0001) and non-ISU (OR=3.63, CI=1.571-8.39, p=0.003). Altogether, hypochromic anaemia, normochromic anaemia and CD4+ T-helper cytopenia are the most prevalent hemocytopenias in HIV infected and uninfected injection and non-injection substance users.

## Introduction

Of the 36.9 million people living with HIV/AIDS in the world, 25.8 million reside in Sub-Saharan Africa (1). In addition, out of the 2 million new HIV infections globally, 1.4 million are recorded in Sub-Saharan Africa (1,2). Substance use has been implicated in the soaring HIV burden and HIV-disease progression in the world (3–20). For instance, the United Nations Office on Drugs and Crime (UNODC) report published in 2018 indicates that 12.5% of Injection Substance Users (ISU) were infected with HIV by the end of 2017 with 5.6% of the global population having used illicit substances (21–25). Injection drug use has been described as one of the major factors that propel the HIV burden worldwide through the practice of sharing hypodermic needles and engaging in unprotected sex with drug and non-drug users (6,26,27) heightening HIV transmission in injection drug users compared to the general population (5). In addition, People-Who-Inject-Drugs (PWID) make up 30% of the new HIV infections in the world (28,29). Non-injection substance use also increases the risk of HIV infection due to altered judgment and increased risky sexual behaviours in non-ISU (9,22,30). Both illicit substance use and HIV infections are increasing in the African continent especially in urban and coastal regions (10,23,24,26,31–38). There are 1.02 million ISUs in Africa out of whom 123,420 are infected with HIV(1,28,29).

HIV infections and illicit substance use have individually been implicated for derangements in the hematologic and immune profiles (5,27,39–49). For instance, HIV infections cause alterations in the hematologic measures. Anaemia, leukopenia and thrombocytopenia are the most frequent hematologic manifestation in HIV-infected individuals (46,50,51). In addition, CD4^+^ T-helper lymphocyte counts are decreased in HIV infection; with lower CD4-counts exhibited in HIV-infected persons compared to HIV-uninfected individuals. (52). Immune status is routinely based on CD4+ T-cell counts (53) Likewise, substance use has been associated with haematological and immune perturbations in HIV uninfected injection and non-injection substance users (26,54). For instance, neutrophilia has been observed in heroin and opium addicts (43,55) while neutropenia, eosinopenia and lymphopenia have been associated with the abuse of Marijuana (*Cannabis sativa*) (56) and chronic alcoholism (57). Monocytosis has been observed in individuals who use Khat (*Catha edulis*) (58) while monocytopenia associated with *Cannabis* use and alcoholism hence decreased proliferation and impaired monocyte and macrophage function (56,57). It is, therefore, possible that these derangements in the haematologic and immune profiles are exacerbated in HIV-positive substance users. (46,50,51). Immune status is an important marker of HIV disease progression and a strong determinant for the initiation of therapy (59). Thus, assessment of anaemia levels and cellular morphology is important in elucidating the underlying mechanisms associated with these observed blood derangements which will, in turn, support the treatment of drug use by delivering services aimed at reducing the adverse health consequences of substance use. Malnutrition has been observed in both HIV-infected and HIV-uninfected injection substance users (4). Irregular carbohydrate, lipid and protein metabolism have been documented in heroin, crack-cocaine addicts and cigarette smokers (60). HIV-1 viral load in-conjunction with immunosuppression have been utilized as markers of HIV disease progression and the initiation of antiretroviral therapy treatment (59,61–65). HIV-1 disease progression has been observed to increase in heroin abusers (4,50,66). Routine hematologic and immune status evaluations guide disease classification and quality management of patients. However, the interplay between HIV infection and substance use on hematologic profiles has not been reported among substance users in Kenya. This study investigated erythrocyte measures, anaemia (levels, types and aetiology) and its association with under-nutrition, immunosuppression and viral failure in Kenyan illicit substance users.

## Materials and methods

### Study site, design and population

This cross-sectional immune-hematologic study was conducted among HIV-1-positive and HIV-1-negative injection substance (ISU) and non-injection substance (non-ISU) users in Mombasa, a coastal Kenyan city. All HIV-1-positive participants in this study had not been previously initiated on any antiretroviral treatment regimen. The detailed description of the study site and study population are published elsewhere (30). The study population was stratified as follows: 1). HIV-positive Injection substance users (HIV+ISU+); 2). HIV-negative Injection Substance Users (HIV−ISU+); 3). HIV-positive non-injection substance users (HIV+ISU-); 4) HIV-negative non-injection substance users (HIV−ISU-) and 5). controls, who never consumed any of the illicit substances as described in the UNODC registry (28,29).

### Ethical considerations

Ethical approvals for the study was obtained from the Kenyatta University (Protocol KU/R/COMM/51/32-4) and the Masinde Muliro University of Science and Technology (Protocol MMU/COR:403012-vol2[8]) institutional review board (IRB). All the respondents were exhaustively educated as per the recommended guidelines (67) and written informed consent obtained prior to enrollment.

### Body mass index (BMI)

Anthropometric measures were obtained from each study participants at enrolment as per the Centres of Disease Control guidelines (68). Height (m) was measured to the nearest 0.1 cm using the Health-o-meter PORTROD wall mounted height rod (Health O meter®, McCook, USA). Study participants were weighed in kilograms (kg) using a portable digital weight scale (Richforth Electronics Co., Fuzhou, China). The BMI was calculated using the height and weight measurements as previously described (68) and BMI<18.5 Kg/m2 defined as underweight.

### Collection of blood samples

5ml of venous blood samples was collected from the freely consenting participants by a certified phlebotomist using a vacutainer assembly into two EDTA and Serum Separating Tubes (SST), BD vacutainer™ tubes (BD, Franklin Lakes, USA). Blood was collected between 8.00am and 10.00am prior to the participants having breakfast to control for the haematological changes due to the circadian rhythm and nutritional status hence obtaining strictly comparable values. All laboratory tests were performed within two hours of sample collection to maintain sample integrity. EDTA blood was used for haematological analysis while SST was used for serum extraction in HIV-1 viral load quantification.

### Hematologic measurements

Complete Blood Counts were done within the first hour of blood collection using the quantitative BC-3200 Mindray auto-haematology analyser (Mindray™ Inc., Mahwah, USA). Anaemia levels and types were classified based on haemoglobin concentration prescribed by the World Health Organization (41) while anaemia aetiology was classified based on blood-markers, cellular morphology and staining characteristics (54–56).

### Preparation of blood slides

Thin blood films were made on new microscope slides (labelled with participant ID) to prevent cell aggregation and stain precipitation. Back up smears were also made. The thin smears were thoroughly air-dried followed by methanol fixation for 10 minutes. The blood smear was then completely covered with undiluted Leishman Stain which was added dropwise using a bulb-pipette. Twice the volume of buffered water (pH. 6.8) was gently added and thoroughly mixed. Staining was done for 10 minutes after which the slide was washed off under running tap water. The back of the slide was wiped and the slide placed standing on a draining rack for the smear to dry.

### Microscopic analysis

Examination of the stained blood films was done by two independent and blinded hemato-technologists who assessed erythrocyte morphology. Slides with differences of more than 5% in the results of the two hemato-technologists were re-read by a third independent hemato-technologist. Ten per cent (n=55) of the read slides were randomly selected and the results confirmed by a haemato-pathologist.

### CD4+ T-cell enumeration

Fifty microlitres (50μl) of EDTA anticoagulated blood was stained with anti-CD3 fluorescein isothiocyanate (FITC), anti-CD4 phycoerythrin (PE) and anti-CD45 peridinin chlorophyll protein (PerCP) fluorescent-labelled mouse-anti-human monoclonal antibodies (BD Tri-test Kit™) (62). CD4+ T cell counts were determined using a BD FACSCalibur flow cytometer (Becton-DickinsonTM, Franklin Lakes, USA). CD4+− T-helper cell counts <500 cells/μ/ was defined as immunosuppression (69).

### HIV-1 viral load determination

RNA was extracted from 200 μl of serum in accordance with the Abbott m2000sp sample preparation system protocol. HIV-1 viral loads were then determined using the automated Abbott m2000SP Real-Time System according to the manufacturer’s instructions (Abbott Molecular Inc., Illinois, U.S.A). The lower limit of viral load quantification was 150 (2.18 log_10_) copies/mL of serum. Virological failure as defined as HIV-1 viral load ≥1000 copies/mL (70).

### Statistical analysis

Statistical analysis was done in RStudio Version 1.1.383 (©2009-2017 RStudio, Inc.) The continuous variables such as weight, height and BMI that were normally distributed were compared across the groups using a one-way ANOVA test. Absolute CD4 counts and erythrocytic measures were compared using non-parametric ANOVA (Kruskal-Wallis Test) followed by Bonferroni post-hoc corrections for multiple comparisons. Viral-loads were compared between the two groups using the Mann-Whitney test. Binary logistic regression analysis was performed within each group to examine the association of anaemia, with under-nutrition, immune-suppression and HIV-1 viral failure; controlling for age, gender, duration and frequency of substance use in injection substance users while age and gender were controlled in the non-injection substance users and controls. All tests were two-tailed and p values <0.05 were considered statistically significant.

## Results

### Anthropometric measurements, CD4 and viral load

Demographic measures, CD4 counts and viral load are as presented in Table 1. A total of 550 adults (males, n=355 and females, n=195) were recruited into the study. This comprised of HIV-positive injection substance users (HIV+ISU+, n=62), HIV-negative injection substance users (HIV-ISU+, n=213), HIV-positive non-injection substance users (HIV+ISU-, n=33), HIV-negative non-injection substance users (HIV-ISU-, n=186) and non-substance using controls (n=56). The median age (years) was significantly different among the study groups (p=0.0047) with posthoc analysis indicating higher median age in HIV+ISU- (p=0.0499) and HIV-ISU- (p=0.0112) compared to the controls. Median height (m) was different across the study groups (p<0.0001) such that HIV-ISU+ were taller than the HIV+ISU- (p=0.0061) and HIV-ISU- (p<0.0001). Similarly, weight (kg) differed across the study groups (p<0.0001) and was higher in HIV+ISU+ than HIV-ISU- (p<0.0001). In addition, under-nutrition rates were higher in HIV+ISU+ (32.3%), HIV-ISU+ (47.4%). HIV+ISU- (48.5%) and HIV-ISU- (22.6%) compared to the healthy controls (8.9%).

CD4 T-helper cell counts varied across the groups (p<0.0001). These were depressed in HIV+ISU+ (median=519 cells/μl, IQR=471) compared to HIV-ISU+ (median=905 cells/μl, IQR=639, p<0.0001), HIV-ISU- (median=859 cells/μl, IQR=515, p<0.0001) and healthy controls (median=774 cells/μl, IQR=461, p=0.0014). Moreover, immunosuppression was most prevalent in HIV+ISU+ (46.8%), HIV-ISU+ (22.1%), HIV+ISU- (36.4%) compared to the HIV-ISU- (16.7%) and controls (17.9%).

HIV-1 viral copies were higher in HIV+ISU+ (median=344copies/μl) compared to HIV+ISU- (median=150copies/μl) However, these differences were not statistically significant. The rates of viral failure (≥1000 copies/μl) were pronounced in both groups (HIV+ISU+, 53.2% and HIV+ISU- (54.5%).

**Table 1.**
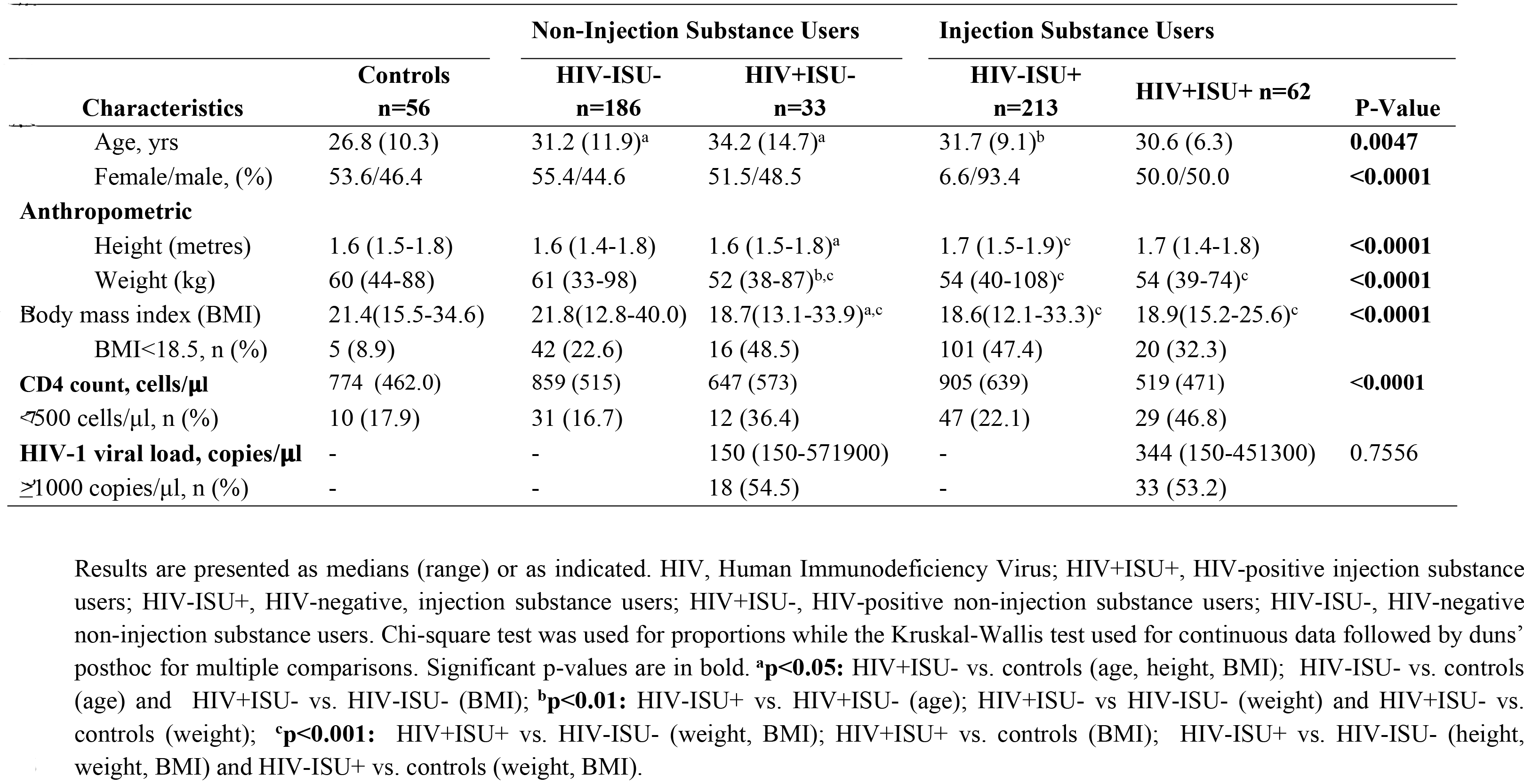
Anthropometric measures, CD4 and viral loads.

### Erythrocyte measures

Erythrocyte measures are summarised in Table 2. The median erythrocyte counts differed across the groups (p=0.0029) with higher counts in the HIV-ISU+ (median, 4.9×1012/L; IQR=0.2) relative to HIV+ISU- (median, 4.0×1012/L; IQR=1.2; p=0.0078). Haemoglobin concentration also differed amongst the groups (p<0.0001), and was elevated in the HIV- ISU+ (median, 12.6g/dL; IQR=2.3) compared to the HIV+ISU- (median, 11.6g/dL; IQR=3.3; p=0.0077). Similarly, HIV+ISU- had lower haemoglobin compared to the HIV-ISU- (median, 12.8g/dL; IQR=2.5; p=0.0003) and controls (median, 13.5g/dL; IQR=2.7; p<0.0001). Moreover, haematocrit was altered across the groups (p=0.0025) and was raised in HIV-ISU+ (median, 41.8%; IQR=6.4) relative to HIV+ISU- (median, 38.5%; IQR=11.7; p=0.0379) and HIV-ISU- (median, 39.7%; IQR=7.8; p=0.0073).

The median mean corpuscular volume (MCV) was not similar across the study groups. The mean corpuscular haemoglobin (MCH) values differed across the study groups (p=0.0020). Depressed MCH levels were observed in HIV-ISU+ (median, 26.4pg; IQR=3.7) compared to the controls (median, 29.0pg; IQR=52; p=0.0006). Meanwhile, the mean corpuscular haemoglobin concentration (MCHC) differed amongst the groups (p<0.0001) and was low in HIV-ISU+ (median, 31.5g/dL; IQR=4.2) relative to HIV-ISU- (median, 31.9g/dL; IQR=2.7; p<0.0001) and controls (median, 32.5g/dL; IQR=2.8; p=0.0004). The red cell distribution width (RDW) was different across the groups (p=0.0036) with posr-hoc analysis indicating lower levels in HIV-ISU+ (median,13.5%; IQR=2.3) in comparison to HIV-ISU- (median, 14.7%; IQR=3.2;, p=0.0029).

**Table 2.**
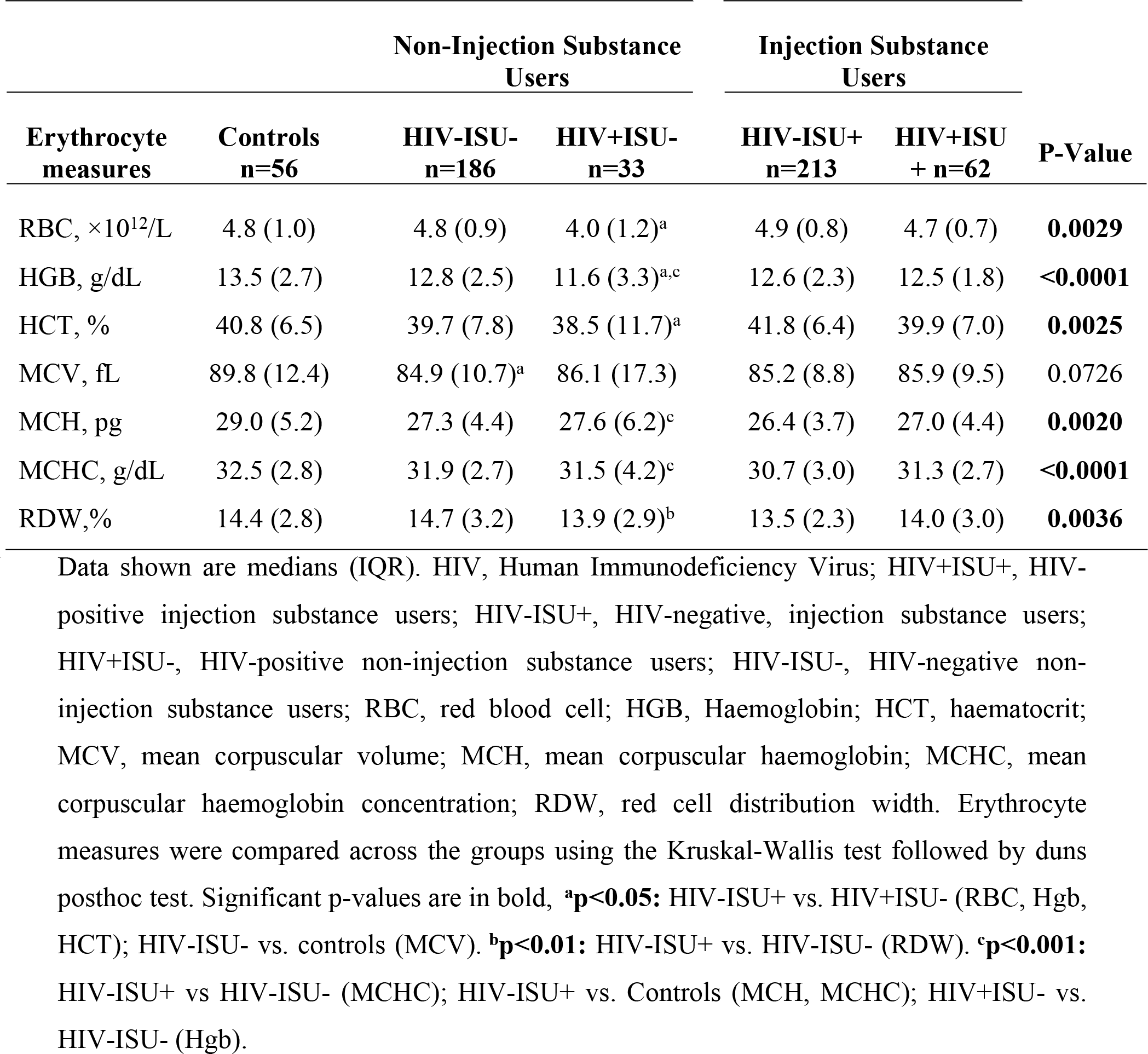
Erythrocyte measures.

### Anaemia levels, types and aetiology

The overall rates of anaemia were higher in HIV-positive subjects (ISU, 48.4% and non-ISU, 63.6%) and HIV-negative ISU (56.3%) relative to the HIV negative non-ISU (39.2%) and controls (28.6%) (Fig 1). Most of the anaemia was mild and moderate (HIV-positive ISU, 56.7% and 40%, and non-ISU, 33% and 52.4%; HIV-negative ISU, 66.7% and 32.5%, and non-ISU, 63% and 28.8%), respectively.

**Fig 1.**
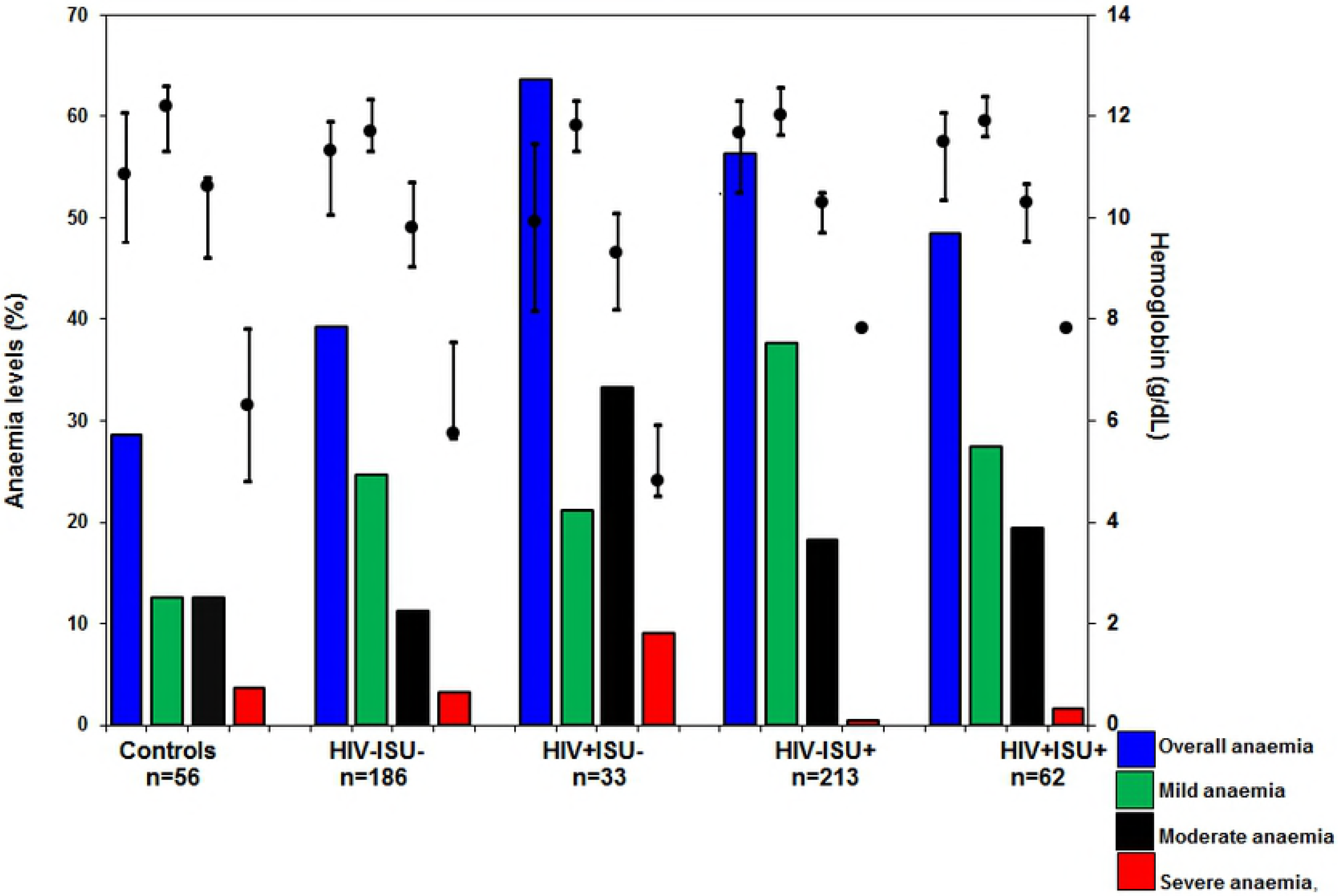
Anaemia levels across the study groups. Primary axis shows anaemia levels represented by the shaded bars. Secondary axis shows the haemoglobin concentration where the whiskers (−) represent the 25^th^ and 75th percentiles for haemoglobin values while the dot (·) represent the median haemoglobin value. p-values are for the haemoglobin concentration within each anaemia level. HIV-ISU-, HIV negative non-injection substance user; HIV+ISU-, HIV positive non-injection substance users; HIV-ISU+, HIV negative injection substance users; HIV+ISU+, HIV positive injection substance users.

Based on RBC morphology, the most prevalent anaemia was hypochromic and normochromic anaemia: (HIV-positive ISU, 50% and 46.7%; and non-ISU, 61.9% and 38.1%; HIV-negative ISU, 63.3% and 35%, and non-ISU 39.7% and 60.3%). Hyperchromic anaemia was less common manifesting amongst the HIV+ISU+ (3.3%), the HIV-ISU+ (1.7%) and controls (6.3%) (Fig 2).

**Fig 2.**
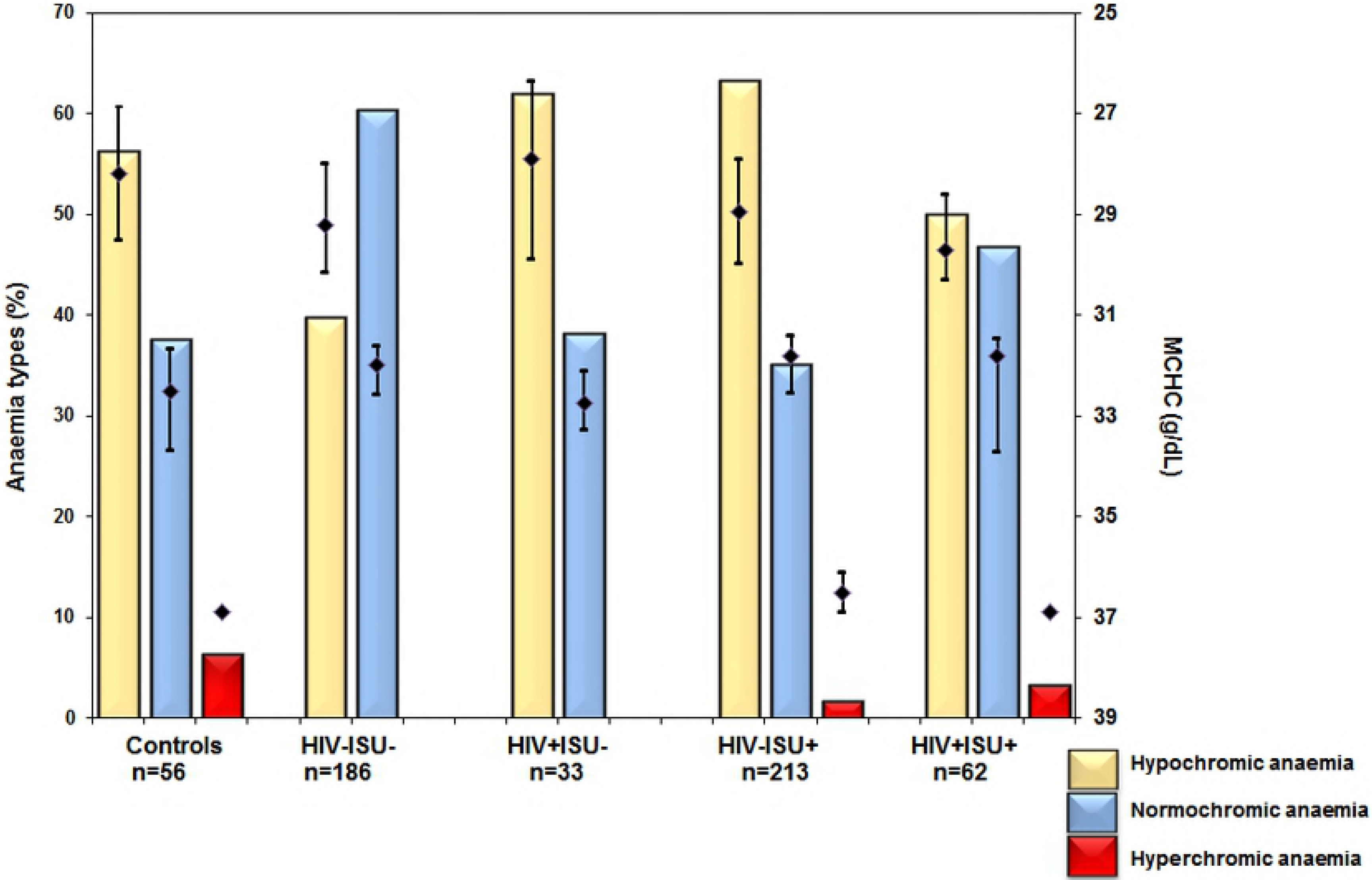
Anaemia types across the study groups. Primary axis shows anaemia types represented by the shaded bars. Secondary axis shows the mean corpuscular haemoglobin concentration (MCHC) where the whiskers (−) represent the 25^th^ and 75th percentiles for MCHC while the dot (·) represent the median. p-values are for the MCHC values within each anaemia type. HIV-ISU-, HIV negative non-injection substance user; HIV+ISU-, HIV positive non-injection substance users; HIV-ISU+, HIV negative injection substance users; HIV+ISU+, HIV positive injection substance users

Anaemia due to mixed aetiology was the most prevalent (49.6%), followed by chronic inflammation (22.9%), nutritional deficiency (15.8%) and blood loss (12.7%) (Fig 3).

**Fig 3.**
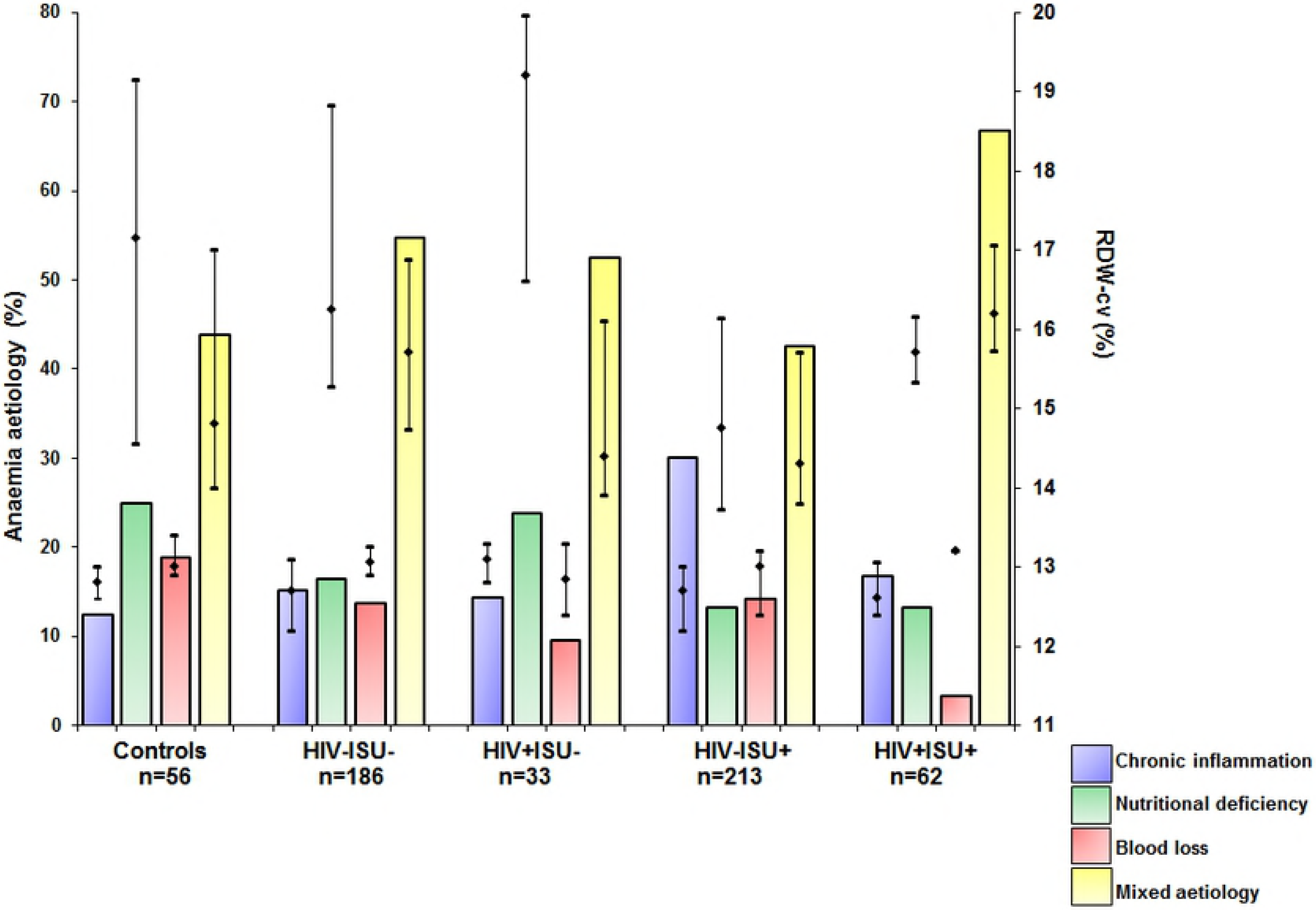
Anaemia aetiology across the study groups. Primary axis shows anaemia aetiology frequency as represented by the shaded bars. Secondary axis shows the red cell distribution width (RDW) where the whiskers (−) represent the 25^th^ and 75th percentiles for RDW while the dot (·) represent the median. p-values are for the RDW values within each aetiology. HIV-ISU-, HIV negative non-injection substance user; HIV+ISU-, HIV positive non-injection substance users; HIV-ISU+, HIV negative injection substance users; HIV+ISU+, HIV positive injection substance users

### Association of anaemia with undernutrition, immune suppression and viral failure

Regression analysis within the HIV+ISU+ and HIV+ISU- indicated that anaemia was neither associated with under-nutrition, immunosuppression or viral failure. However, anaemia was associated with immunosuppression, amongst the HIV-ISU+, (OR=3.952, CI=1.934-8.077, p<0.0001) and HIV-ISU- (OR=3.630, CI=1.571-8.390, p=0.003).

## Discussion

Anaemia is characterized by the insufficiency in the number of red blood cells, consequently affecting their oxygen carrying and delivery capacity to tissues (71). Anaemia in Human Immunodeficiency Virus (HIV) infected persons is life threatening as it is associated with enhanced HIV disease progression hence diminished survival (72). Substance use, on the other hand, has been associated with varied haematological derangements including anaemia (73–75). This cross-sectional study investigated the interplay between substance use, HIV infection and anaemia in Kenyan injection and non-injection substance users not under any active antiretroviral treatment. Erythrocyte indices, anaemia levels, type and aetiology were determined. It was observed that HIV-negative illicit substance users with drug-induced-immune-suppression were thrice as likely to develop anaemia compared to their HIV-positive counterparts. This is important in fostering the treatment and management of illicit substance users while reducing the adverse health consequences of substance use.

The overall prevalence of anaemia was highest amongst the HIV-positive non-injection substance users, HIV negative injection substance users and HIV positive injection substance users. Typically, anaemia was more severe in HIV positive substance users compared to HIV negative substance users. It is likely that HIV-infection in substance users aggravates anaemia. HIV has been shown to replicate in other cells of the haematopoietic lineage other than the immune cells thus leading to the haematological derangements, with erythroid dysplasia observed as a common feature upon bone marrow examination of people who are infected with HIV (76–78). The overall prevalence of anaemia amongst the healthy controls (28.6%) in the study area of the coastal city of Mombasa Kenya was lower than that of the global prevalence described elsewhere (79), but relatively higher compared to the World Health Organization estimates for the prevalence of anaemia (24.8%) in the general population of Kenya (80). The controls in this study were recruited from asymptomatic individuals within the community. Thus, it was observed that in as much as individuals within a community would seem healthy due to lack of clinical symptoms, laboratory investigations seem to suggest otherwise. High rates of anaemia in the general population from the coastal city are attributable to the extravagant prevalence of malnutrition, chronic protozoal and helminthic infections (81–83).

Generally, anaemia rates were high in the substance-using groups compared to the controls suggesting that illicit substance use is associated with anaemia which is exacerbated by HIV infection. It is possible that drug metabolites negatively influence erythropoietic hormones and may trigger intravascular haemolysis and premature splenic destruction of red blood cells. However, this hypothesis needs to be substantiated with further research on the same. Mild and moderate anaemia were the most prevalent types of anaemia based on haemoglobin concentration (71). Severe anaemia was the least recorded type of anaemia as individuals in this state are either bedridden or comatose. However, we were able to observe few cases of individuals with severe anaemia who were neither comatose nor bedridden across all the study groups. A possible explanation is that the anaemia amongst these individuals might have developed over long periods of time providing room for the physiologic compensatory mechanisms to kick in hence allowing greater loss of red blood cell (RBC) mass over time without any obvious clinical symptoms (84).

Chronic inflammation was the second most common mechanism associated with anaemia prevailing in injection and non-injection substance users. Therefore, substance use is likely to be associated with inflammation. Khat and alcohol use has been shown to cause intestinal lesions leading to gastritis (85–90). This intestinal inflammation is likely to cause the liver to secrete more of the hormone hepcidin which acts by preventing the body from utilizing stored iron (ferritin) and subduing iron absorption in the duodenum. As a matter of fact, anaemia due to nutritional deficiency was the third most common cause across all the study participants. Nutritional deficiency anaemia is probably due to the low dietary intake of iron, folate and vitamin B12 in the general population and substance-induced damage of the gastrointestinal mucosa within the substance using groups (91). Mal-absorption states in these groups need to be investigated including the production and inhibition of the intrinsic factor, which is important in differentiating the types of nutritional anaemias.

Anaemia due to mixed aetiology was the most frequent mechanism across our study participants. However, due to the limited resources and time constraints, we could not perform further investigations to specifically determine the kinetics underlying the mixed aetiology of anaemia. Despite this challenge, reports from our analysis indicated a coexistence of the above mechanisms with other aetiologies whose haematological “blueprints” were suggestive of underlying hemoglobinopathies and thalassemias. However, this claim needs to be substantiated by further investigations. In addition, there were wispy signs indicative of intravascular haemolysis and suppression of erythropoiesis. We speculate that intravascular haemolysis could be attributable to the damping effect where the drug metabolites are adsorbed onto the RBCs which become antigenic resulting in their untimely destruction by the immune and the reticuloendothelial system.

Anaemia observed was also classified based on the RBC chromasia as hyperchromic, hypochromic and normochromic. Hypochromic anaemia was the most prevalent type of anaemia accounting for more than 50% of the anaemia. Hypochromic anaemia was common across all the study groups. Some of the mechanisms driving the existence of hypochromic anaemia include iron deficiency, toxic anaemia, sideroblastic anaemia, myelodysplasia, haemolytic thalassemia and megaloblastic anaemia (92). Chronic alcohol users have been shown to present with clinical findings suggestive sideroblastic and megaloblastic anaemia (57). In our study, a nutritional deficiency was the third most common cause of anaemia, which could be attributable to insufficient iron supplementation in the diet. Studies have reported the most common cause of anaemia in resource-limited tropical settings include underlying nutritional deficiencies and endemic parasitic infections (93). Substance addicts have been observed to have altered eating habits such as bypassing meals and fasting in order to prolong the effects of the drugs (94). These addicts usually have limited finances which are mainly spent on sustaining their drug habits hence have a lower dietary intake of fruits, vegetables and other animal products. As such, they are prone to numerous vitamin deficiencies, some of which are necessary for the synthesis of haemoglobin (such as vitamin B12, folate) while others aid in the absorption of iron from the intestines (e.g. vitamin C). Microcytic hypochromic anaemia was the second most common type of hypochromic anaemia amongst the study participants and has been associated with chronic inflammation and thalassemias (84). Consistent with our findings, previous studies showed that the use of illicit injection substance was associated with normocytic hypochromic anaemia whose main aetiology is the iron deficiency (95). However, the physiological and biochemical mechanisms behind the iron deficiency have not been demonstrated warranting further laboratory investigations to delineate between real nutritional deficiency and iron distribution disorders.

Normochromic anaemia was the second most prevalent type of anaemia. Normochromic anaemia has been associated with a number of mechanisms such as short-term blood loss with adequate physiologic reserves, accelerated red blood cell turnover and suppression of red blood cell production when there is adequate iron intake (92). Normocytic normochromic anaemia was most prevalent in HIV-negative non-injection substance users (37%); HIV infected non-injection drug users (33.3%) and HIV infected injection drug users (30%). Since normocytic normochromic anaemia is more predominant in HIV infected individuals, it is concluded that HIV disease accelerates normocytic normochromic anaemia. The aetiology of normocytic normochromic anaemia has been described elsewhere (96) to be as a result of chronic disease, destruction of red blood cells and the disappearance of erythrocyte precursors from the bone marrow; factors which have been well recorded in HIV disease progression (7,97–102).

Haemoglobin levels were significantly lower in HIV positive non-injection drug users compared to controls, HIV naïve non-injection drug users and HIV naïve injection drug users. This observation proposes that HIV infection may be the culprit resulting in reduced haemoglobin. This is backed up by a study which revealed that advanced HIV progression is marked by a reduction in haemoglobin (97) due to alterations in cytokine production affecting other homeostatic processes such as hematopoiesis; autolysis and Vitamin-B12 deficiency due to impaired absorption (103).

Results from this study show a significant decline in erythrocyte counts in HIV infected non-injection substance users compared to HIV-negative injection substance users. HIV-negative non-ISU exhibit higher erythrocyte levels suggesting that HIV infection may play a role in erythrocyte depression. This finding is similar to a study that investigated the role of HIV in anaemia (72,101). Different mechanisms have been conjectured by which HIV suppress RBC counts and they include marred division and endurance of hematopoietic progenitor cells (98,99), aberrant cytokine production such as erythropoietin by stromal cells and autoimmune responses resulting in the untimely destruction of red blood cells in the spleen and by autoantibodies (101).

On the other hand, the relatively normal erythrocyte counts in HIV-infected injection substance users similar to that of the controls suggest that injection substance use seem to ameliorate RBC populations in HIV infected individuals. This finding is similar to a different study where opium and heroin-dependent individuals did not exhibit significant differences in their erythrocyte populations compared to the healthy groups (43). However; despite the fact that RBC population is not significantly altered in number; erythrocyte function is altered as shown in a different study (104) where the red cell immune-adherence function was significantly decreased in heroin users. Therefore, it would be of great interest to investigate erythrocyte function amongst these groups.

Immune status was classified based on the Centers for Disease control guidelines (105). Results from this study show that immune suppression was marked in HIV positive substance users. This is attributable to the fact that HIV virus replicates in immune cells causing their premature death upon pyroptosis (37,44,101). Immunosuppression was also observed in illicit substance users who were HIV negative. Studies conducted on non-human primates have demonstrated the immunosuppressive effects of morphine on the immune cells (106). In the aforementioned study, T-cell activation in non-human primates was significantly decreased upon morphine administration with negligible changes in T-cell, neutrophil and natural killer cell counts. Proteomic analysis in this study showed a significant decrease in the protein Ki-67+. The Ki-67+ is an important signalling molecule that aids in cellular proliferation (107,108). It would be of very much interest to investigate the proteome and metabolome within our study population to better understand the alterations in their physiological processes.

The regression analysis outcomes of this study suggest that HIV negative substance users with drug-induced immunosuppression are likely to develop anaemia compared to the HIV positive substance users. We speculate that this observed association may be as a result of deficiencies in one or more micronutrients other than iron such as copper and zinc that may be critical for both immune function and production of haemoglobin by modulating enzymes associated with these processes. However, this assertion needs to be tested and substantiated by further studies.

## Conclusion and recommendations

Haematological, immune and nutritional parameters are influenced by infections with HIV and substance use. Combinations of these two factors exacerbate anaemia and other haematological anomalies. Haemoglobin levels and red blood cell indices are significantly altered in HIV infected substance users compared to HIV negative substance users. Examination of the bone marrow for erythroblasts and reticulocyte counts are warranted to determine the effect of substance use and HIV on haematopoiesis in these individuals. In addition, there is the need for further biochemical tests such as serum iron, ferritin, total iron binding capacity (TIBC), transferrin, folate, cobalamin, vitamin-C bilirubin and haptoglobin concentration, including testing for liver enzymes, cytokines and kidney function tests to examine the rate of RBC turnover in these individuals.

## Acknowledgements

We thank the study participants for making this study possible. We are grateful to the management and staff of the Bomu Hospital for their support during the study. This study was supported, in part, by the Kenya National Commission for Science, Technology and Innovation [NCST/5/003/065] grants to TW and VB.

